# Snappy: *de novo* identification of DNA methylation sites based on Oxford Nanopore reads

**DOI:** 10.1101/2025.08.03.668330

**Authors:** Dmitry N Konanov, Danil V Krivonos, Vladislav V Babenko, Elena N Ilina

**Affiliations:** Research Institute for System Biology and Medicine; Lopukhin Federal Research and Clinical Center of Physical-Chemical Medicine of Federal Medical Biological Agency

## Abstract

**Summary:** Nowadays, search for methylation sites in bacteria is usually performed by direct detection of nucleotide motifs over-represented in modified contexts, using classical motif enrichment approaches oriented only on context sequences themselves. Herein, we present a new algorithm Snappy, which is actually rethinking of the original Snapper algorithm but does not use any enrichment heuristics and does not require control sample sequencing. Opposite to previous methods, Snappy uses raw basecalling data simultaneously with the motif enrichment process, thus significantly enhancing the enrichment sensitivity and accuracy compared with other enrichment algorithms. The versatility of the method was shown on both our and external data, representing different bacterial taxa with complex and simple methylome.

**Availability and implementation:** Source code and documentation is hosted on GitHub (https://github.com/DNKonanov/ont-snappy) and Zenodo (zenodo.org/records/16731817). For accessibility, Snappy is installable from PyPi using ‘pip install ont-snappy’ command.

## 1. Introduction

In prokaryotes, DNA methylation is provided by DNA methyltransferases (MTases), special site-specific enzymes (or enzymatic complexes) that modify nucleotide bases in dsDNA in specific nucleotide contexts[1]. The most known modification types are methylation of adenine in 6th position, and methylation of cytosine in 4th or 5th positions, producing 6NmA, 5mC and 4NmC respectively. Identification of genome contexts containing modified bases is primarily provided by sequencing technologies designed to analyze native DNA material, such as PacBio SMRT and Oxford Nanopore[2]. Specifically, the last Oxford Nanopore r10 chemistry and new software significantly improved the quality of methylated bases identification[3], thus allowing the analysis of DNA methylomes *de novo*, without sequencing control samples.

For the organisms for which exact sequences of methylation sites are preliminary known, analysis of DNA methylation is quite trivial, and includes direct consideration of specific genome positions containing target sites, and fraction of reads that bring a methylated base in these positions. On the other hand, while sequences of methylation sites are not known, it is first required to carry out the procedure of motif enrichment. Motif enrichment algorithms currently used for identification of methylation sites such as MEME[4] or STREME[5], rely only on nucleotide sequences bringing modified bases, but do not use raw information provided by basecaller.

Herein, we present Snappy, the first tool that combines motif enrichment with simultaneous analysis of basecalling results. As its predecessor Snapper[6], Snappy is primarily oriented on Oxford Nanopore data, but unlike Snapper, it does not use any heuristics, does not require control sample sequencing, and is significantly easier to run. In this study, we validate Snappy on *Helicobacter pylori* - an organism with the most complex methylome[7], and few other bacteria, and compare it with the MicrobeMod pipeline[8], which actually implements STREME algorithm, but with an additional motif correction procedure.

## 2. Implementation

Snappy is a new tool designed for fast and accurate identification of DNA methylation sites mainly based on Oxford Nanopore sequencing data. The algorithm uses an iterative approach, where each iteration consists of two steps: the first step is formation of a short anchor motif variant (up to 6 bases), and the second step is extending and correction of the anchor variant based on the modification probabilities assigned by basecaller for all genomic contexts satisfying the anchor motif or its close derivatives. The detailed description of the algorithm is available in **Supplementary Data 1**, some comments about Snappy behavior are listed in **Supplementary Data 2**. Since Snappy is a fully-automated tool, it does not require any user control during running, unlike specialized pipelines such as Nanodisco[9].

**Input data** required by Snappy, are a **FASTA-file with the genome** of the analyzed organism, and a **bed-file, generated by ‘modkit pileup’**. The modkit bed-file just summarizes the information from MM and ML fields in bam-files, so actually Snappy is compatible with any sequencing and data processing techniques that could provide MM+ML fields in BAM. Snappy does not have additional algorithm hyperparameters and automatically optimizes the threshold values based on the input data.

The main output of Snappy includes 1) **summary table**, where all identified sites are listed, 2) **results table**, where for each identified site all its genomic occurrences and corresponding methylation levels are listed. Additionally, Snappy generates **two types of plots**, representing methylation level of the sites and their localization. In controversial cases, these plots might be used to ensure the correctness of the inferred motifs. For advanced users, who are interested in a more complex analysis, Snappy saves filtered and extended bed-files used during the enrichment process, and regexp-formatted records for each site so that users could conveniently operate Snappy results using instruments such as Pandas or Polars.

**Figure 1.**
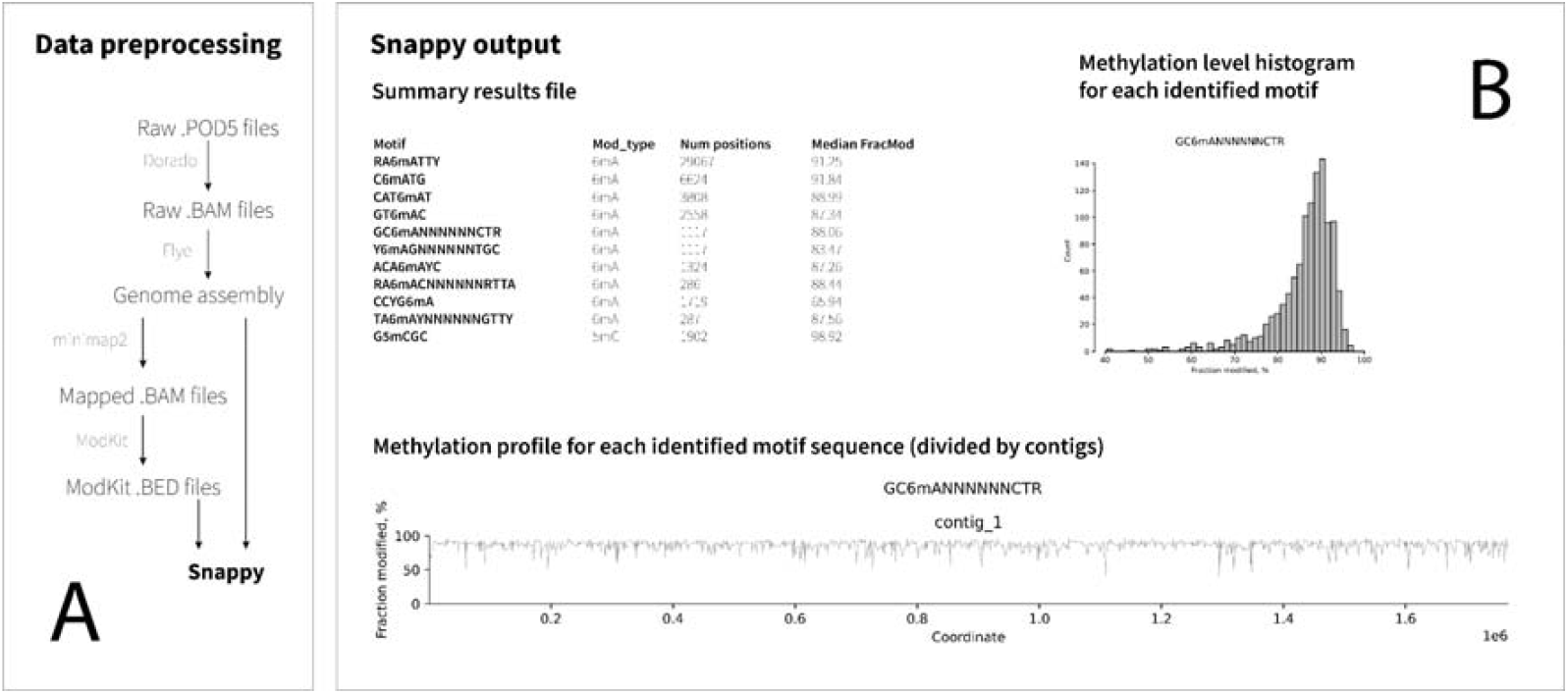
**A)** Data preprocessing procedures required to prepare input files required for Snappy. **B)** The main output provided by Snappy includes both text and graphical files.

## 3. Results

### 3.1. Method validation

The method was first validated on *Helicobacter pylori* J99, which was characterized earlier using both SMRT PacBio and Oxford Nanopore technologies[6,10]. *Helicobacter pylori* is a unique bacterial species with a tremendous variety of restriction-modification systems (up to 30 in some strains), so it has a very complex DNA methylation pattern, with a lot of overlapping or similar methylation sites. Due to that, it was chosen as a validation object. Snappy was compared to the MicrobeMod pipeline, which uses STREME motif enrichment algorithm, but with an additional motif correction procedure.

In result, Snappy outperformed MicrobeMod, demonstrating higher motifs enrichment accuracy for less computational time, and identified all known *H. pylori* J99 motifs. The only motif that was identified incompletely by Snappy was RTAYnnnnnRTAY, because its subvariant GTACnnnnnATAT was not present in the reference genome, so the algorithm could not check if it was modified or not. All other RTAYnnnnnRTAY subvarians (RTAYnnnnnGTAT, RTATnnnnnRTAT, and ATAYnnnnnATAY) were successfully enriched, and the users can decide by themselves if these subvariants should be merged.

**Table 1.**
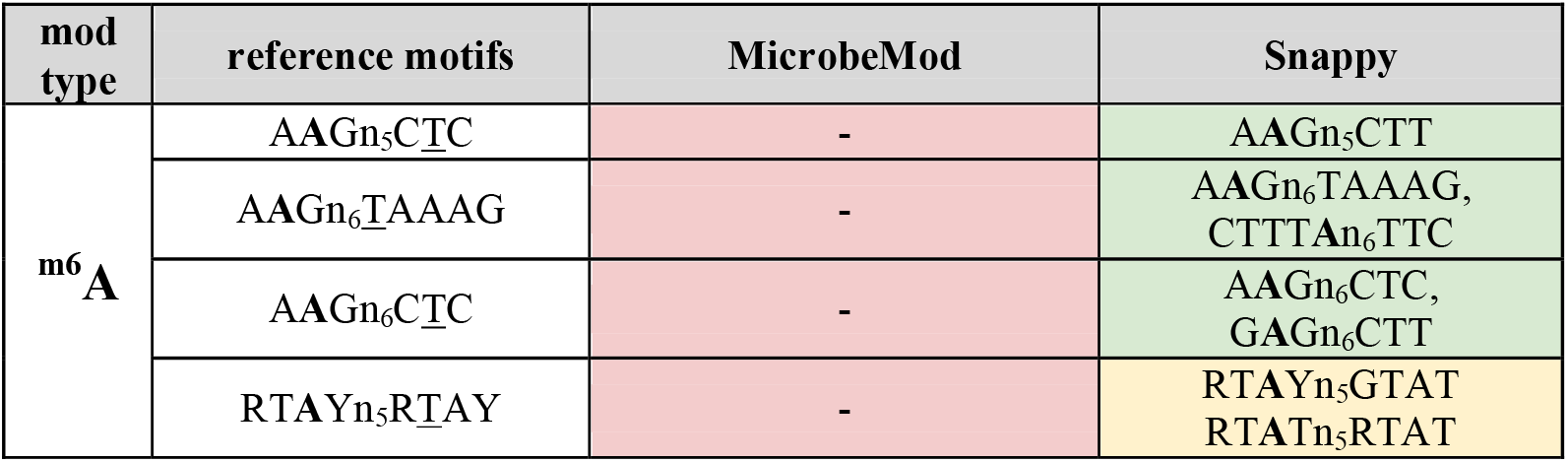
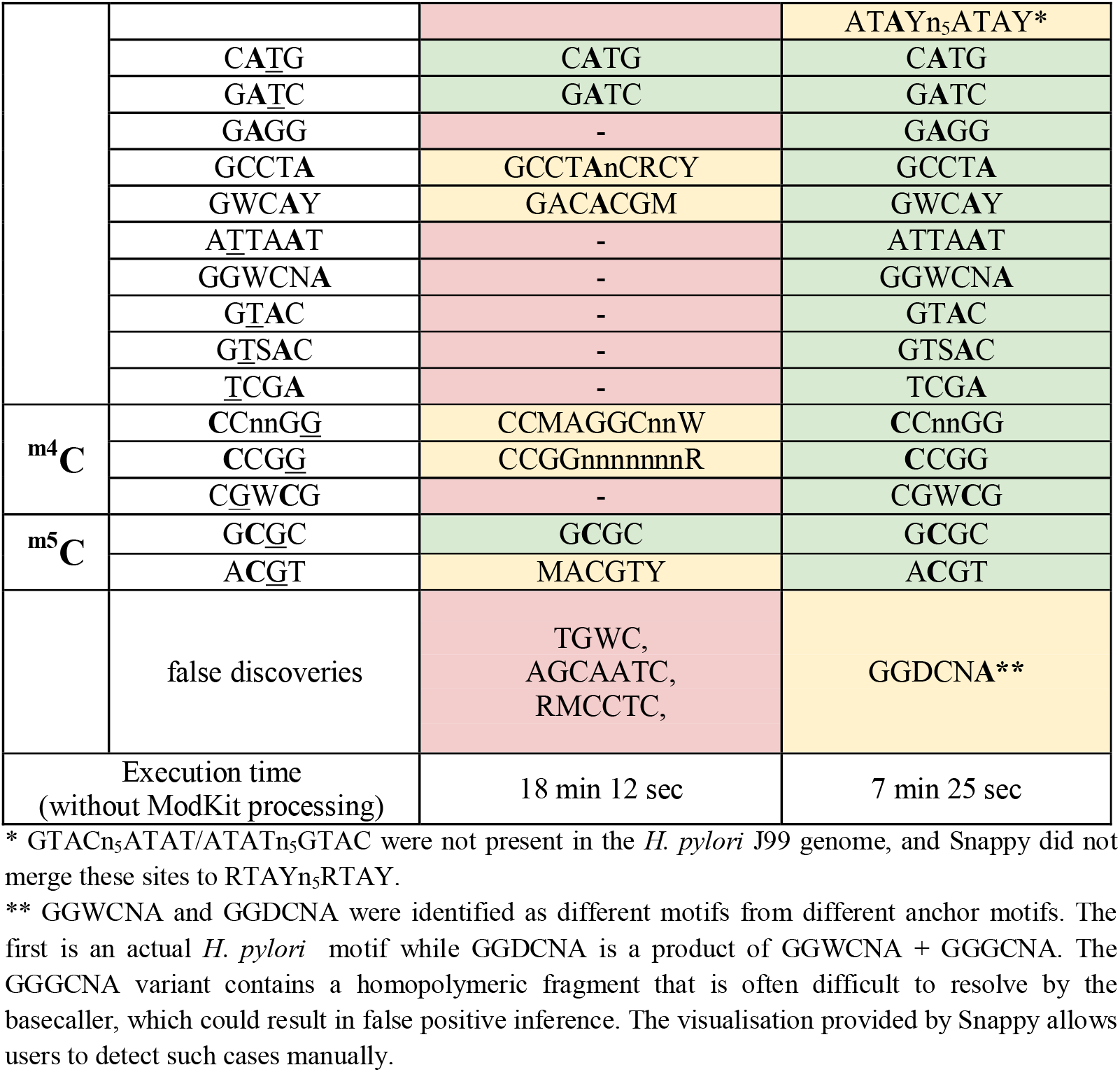
Identification of methylated sites in *H. pylori* J99.

### 3.2. Identification of methylation sites in other bacteria

To demonstrate versatility of the method, we additionally analyzed publicly available Oxford Nanopore sequencing data, for which POD5-files were available. Here, we first chose *Campylobacter* species for the analysis, since their genomes are also known to encode a lot of different R-M systems. Also, to demonstrate how the algorithm works with organisms with a less complex methylome, we ran Snappy on *Escherichia coli* and *Mycobacterium tuberculosis* genome sequencing data. *C. jejuni* ATCC 33560, *C. lari* ATCC 35221, *E*.*coli* ATCC 25922, and *M. tuberculosis* mc26030 raw sequencing data were obtained from The Melbourne University data repository (https://figshare.unimelb.edu.au/) and processed with both Snappy and MicrobeMod.

For *C. jejuni* 33560, all 9 motif sequences identified by Snappy were complete and correct according to REBASE. Seven of them were confirmed using REBASE as well as presence of corresponding DNA MTases in the genome. Two other sites also had records in REBASE for *C. jejuni* but were not linked to corresponding MTase genes (rebase.neb.com/cgi-bin/pacbioget?13650, rebase.neb.com/cgi-bin/pacbioget?42913). MicrobeMod returned only one correct motif sequence CATG, and four incompletely identified motifs.

In *C. lari* 35221, Snappy identified five methylation sites. Four of them were confirmed using REBASE, but GARAnnnnnnnnTAC site identified by Snappy differed from GADAnnnnnnnnTAC site in REBASE by one degenerate base. We manually checked methylation calling results generated by Dorado for GATAnnnnnnnnTAC contexts and did not find a significant methylation level (**Supplementary Figure 1**). For *C. lari*, none of the sites returned by MicrobeMid were identified correctly.

In *M. tuberculosis* mc26030, Snappy identified only one methylation site. According to REBASE, this strain has one more I type R-M system specific to the GCAYnnnnATC site. We manually checked adenine methylation level in genome positions satisfying this site and did not observe any adenine methylation signals (**Supplementary Figure 2**). Probably, although the genes encoding this R-M system are present in the genome, it was not active in that specific sample.

In *E*.*coli* ATCC 25922, Snappy identified five methylation sites. Four of them (GATC, CCWGG, RTACnnnnGTG, and GGTCTC) were in full accordance with REBASE records for this specific strain (tools.neb.com/genomes/view.php?seq_id=90864&list=1). The “Y” degenerate base in the fifth site GAGAYC was a result of superposition of actual *E. coli* methylation motifs GAG**A**CC and G**A**TC, both methylated in adenine. Actually, such collisions could be resolved only experimentally. Additionally, in *E. coli* data we first faced systematic false-positive detection of methylation bases caused by neighbourhood effects: the basecaller assigned quite high methylation probabilities for TC**C**nnCCTGG (in a form of TC^**5m**^**C**nnCCTGG), which was most likely caused by total cytosine methylation in the CCTGG submotif (**Supplementary Figure 3**). This artifact was observed manually during the algorithm development. MicrobeMod identified only two main methylation sites - CCWGG and GATC.

**Table 2.** DNA methylation sites identified in external data with MicrobeMod and Snappy.

| strain | MicrobeMod | Snappy |
| --- | --- | --- |
| *C. jejuni* ATCC 33560 | AAATTT<br>CATG*<br>RATATGnnWnW<br>GACAATCA<br>WGCGC | RAATTY<sup>1</sup><br>CATG<sup>1</sup><br>CATAT<sup>2</sup><br>GTAC<sup>1</sup><br>GCAnnnnnnCTR<sup>1</sup><br>ACAAYC<sup>2</sup><br>RAACnnnnnnRTTA<sup>1</sup><br>CCYGA<sup>1</sup><br>GCGC<sup>1</sup> |
| *C. lari* ATCC 35221 | AAATTT<br>CYATNKnnnTTGY<br>CMGGATCA<br>CAAGTnnTWTAGMA<br>ANAGCTGATCAA | RAATTY<sup>1</sup><br>CYAnnnnnnTTG<sup>1</sup><br>GATC<sup>1</sup><br>GARAnnnnnnnnTAC<sup>3</sup><br>GTRGAG<sup>2</sup> |
| *E. coli* ATCC 25922 | GATC<br>CCWGG | GATC<sup>1</sup><br>CCWGG<sup>1</sup><br>RTACnnnnGTG<sup>1</sup><br>GAGAYC<sup>4</sup><br>GGTCTC<sup>1</sup> |
| *M. tuberculosis* mc26030 | CTCCAGNNNSNNS | CTGGAG<sup>1</sup> |

<sup>1</sup> Correctness of the sequences and presence of corresponding MTase genes in the genome were confirmed using REBASE.
<sup>2</sup> These sites have been experimentally confirmed for the studied *Campylobacter* species using PacBio, but corresponding MTases are unknown (rebase.neb.com/cgi-bin/pacbioget?13650, rebase.neb.com/cgi-bin/pacbioget?42913).
<sup>3</sup> This site is listed in REBASE as GADAnnnnnnnnTAC. We manually checked the methylation level of the GADAnnnnnnnnTAC subvariant, and did not observe a significant shift in frac\_mod values (**Supplementary Figure 1**).
<sup>4</sup> “Y” degenerate base in this site is most likely an artifact, since its GAGATC subvariant contains a totally methylated GATC motif. Interestingly, its GAGACC subvariant methylated in adenine is a full complement for the GGTCTC motif methylated in cytosine. Non-symmetric methylation of GGTCTC/GAGACC has been shown earlier for *E. coli* strains[11].

### 3.3. Main algorithm limitations

1. If a methylation motif has subvariants that are not presented in the considered genome, it could be extracted incompletely. Thus, *H. pylori* J99 actually has RTAY…..RTAY methylation site, but its subvariant GTAC……ATAT is not presented in the genome, so the algorithm cannot check if it methylated or not, and returns as a result only RTAY…..GTAT, RTAT…..RTAT, and .ATAY…..ATAY subvariants. The users could decide themselves if they should merge these subvariants.1)
2. If a site is partially methylated, it could not provide satisfactory modality metric value, and could not be enriched by Snappy. In general, such cases are rather rare, and users are recommended to use additional enrichment methods like STREME if they suspect incomplete methylation.
3. If a site has very few occurrences in the genome (typically less than 20) it cannot be identified by Snappy.

## 4. Methods

### 4.1. Acquisition of data used in the study

Genome sequencing of the *Helicobacter pylori* J99 genome was conducted during this study on the Oxford Nanopore platform (MinION) using an R10.4.1 flow cell (FLO-MIN114). *Escherichia coli, Mycobacterium tuberculosis*, and *Campylobacter* species sequencing data in the POD5 format were downloaded from the data repository of The University of Melbourne (https://figshare.unimelb.edu.au/).

### 4.2. Raw data preprocessing

First, POD5 files were processed by Dorado v0.7.1, with additional usage of models for 6mA, 5mC, and 4mC (model versions v5.0). The resulting BAM files were converted to FASTQ using samtools (v1.19.2). FASTQ files were used for genome assembly with Flye (v2.8.1-b1676)^1^. Using minimap2 (2.26-r1175), raw BAM files were remapped to the assembled genome to obtain sorted mapped BAM files with MM and ML fields. These files were processed by Modkit (v0.2.4) with parameter --filter-threshold 0.66 to obtain bed-files next passed to Snappy.

It should be noted that although in this study all datasets were analyzed using --filter-threshold 0.66 regardless modification type, in some cases Modkit parameters should be preliminary optimized for different modification types independently. Thus, considering the total distribution of modification probabilities assigned by Dorado for some modification type, it is recommended to use as a cutoff value the 10th percentile of this distribution to accurately detect this specific modification (https://github.com/nanoporetech/modkit/issues/245). The Modkit tool has a specific ‘sample-probs’ mode that allows users to perform such parameters tuning.

## 5. Discussion and conclusion

The last r10.4.1 version of ONT flow cell and new basecalling models significantly increase accuracy of methylated bases detection and simplify the analysis. Despite that, algorithms currently used for identification of methylation sites continue to rely only on nucleotide sequences surrounding modified bases but do not use low-level information from basecalling results. In our knowledge, Snappy presented in this study is the first attempt to combine enrichment algorithms with simultaneous analysis of raw modification probabilities provided by basecaller Dorado.

Snappy successfully identified all methylation sites in *Helicobacter pylori* and two *Campylobacter* species, thus demonstrating high accuracy in motif extraction on the organisms with a complex methylome. DNA methylation sites in bacteria species with fewer R-M systems such as *Escherichia coli* and *Mycobacterium tuberculosis* were successfully identified as well.

Compared to Snapper designed to work with r9.4.1 flow cell data, Snappy does not use any enrichment heuristics, and does not require control sample sequencing, which, coupled with a new graph-based enrichment algorithm, provides very high processing speed. Moreover, due to a new logic, Snappy enrichs long bipartite motifs as easily as short motifs and uses the same confidence metrics for all motifs, while Snapper runs an additional module for enrichment of long motifs, with usage of special heuristics.

Snappy allowed us to reveal two types of artifacts arising on the basecalling stage, which could affect DNA methylation analysis. First, we faced the neighbourhood effect when positions very close to modified bases were also systematically identified by Dorado as modified. Second, although new ONT chemistry significantly improves resolution of homopolymers, we found that short homopolymers could lead to false positive inference of modified positions. Thus, Snappy has highlighted the ways in which current non-canonial Dorado models can be improved.

## Author contributions

**Dmitry N. Konanov**: Conceptualization, Software, Validation, Visualization, Writing - original draft; **Danil V. Krivonos**: Software, Validation, Visualization, Writing - original draft; **Vladislav V Babenko**: Investigation, Resources, Writing - review and editing, **Elena N. Ilina**: Supervision, Writing - review and editing.

## Funding

This work was supported by the government task No 122030900069-4.

## Conflict of interests

None declared.

## Footnotes

1 https://www.nature.com/articles/s41587-019-0072-8

## Notes

### Competing Interest Statement

The authors have declared no competing interest.

