## Supplementary Materials for "Snappy: *de novo* identification of DNA methylation sites based on Oxford Nanopore reads"

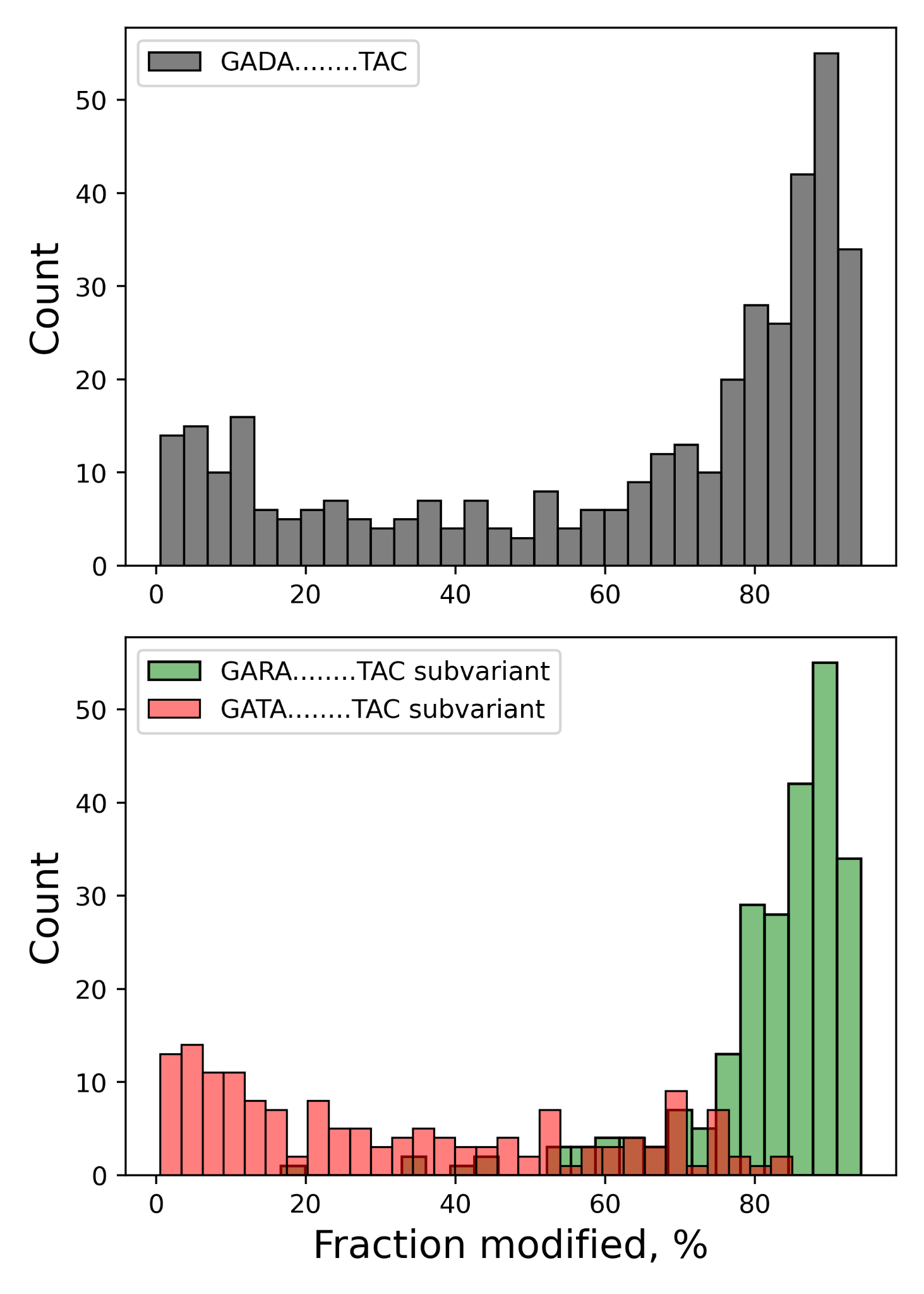

**Supplementary Figure 1.** Manual check of GARAnnnnnnnnTAC site correctness. In REBASE, this site has a form of GADAnnnnnnnnTAC (where D = A/G/T, and R = A/G). The grey histogram indicates that site GADAnnnnnnnnTAC has a visible mode close to zero. The bottom histograms indicate that this mode is explained by the subvariant GATAnnnnnnnnTAC, while the GARAnnnnnnnnTAC subvariant provides a unimodal distribution close to 1.0, so it is totally methylated. Thus, the GARAnnnnnnnnTAC motif identified by Snappy seems to be correct.

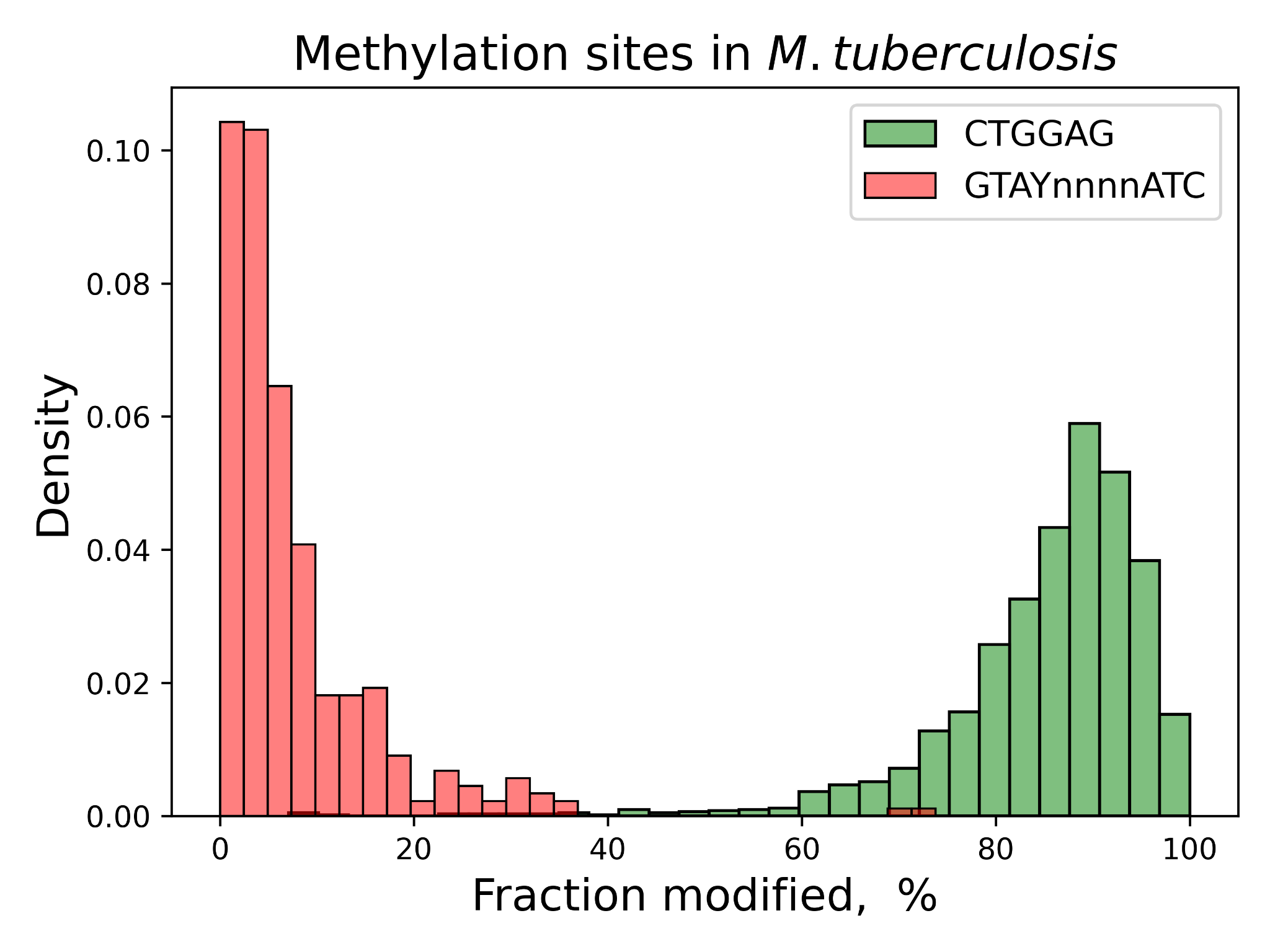

**Supplementary Figure 2.** “Fraction modified" distributions for methylation sites in *M. tuberculosis.* According to REBASE, this strain should have two methylation sites - CTGGAG and GTAYnnnnATC. Snappy identified only the CTGGAG site. Manual check of GTAYnnnnATC confirmed absence of methylation in this site.

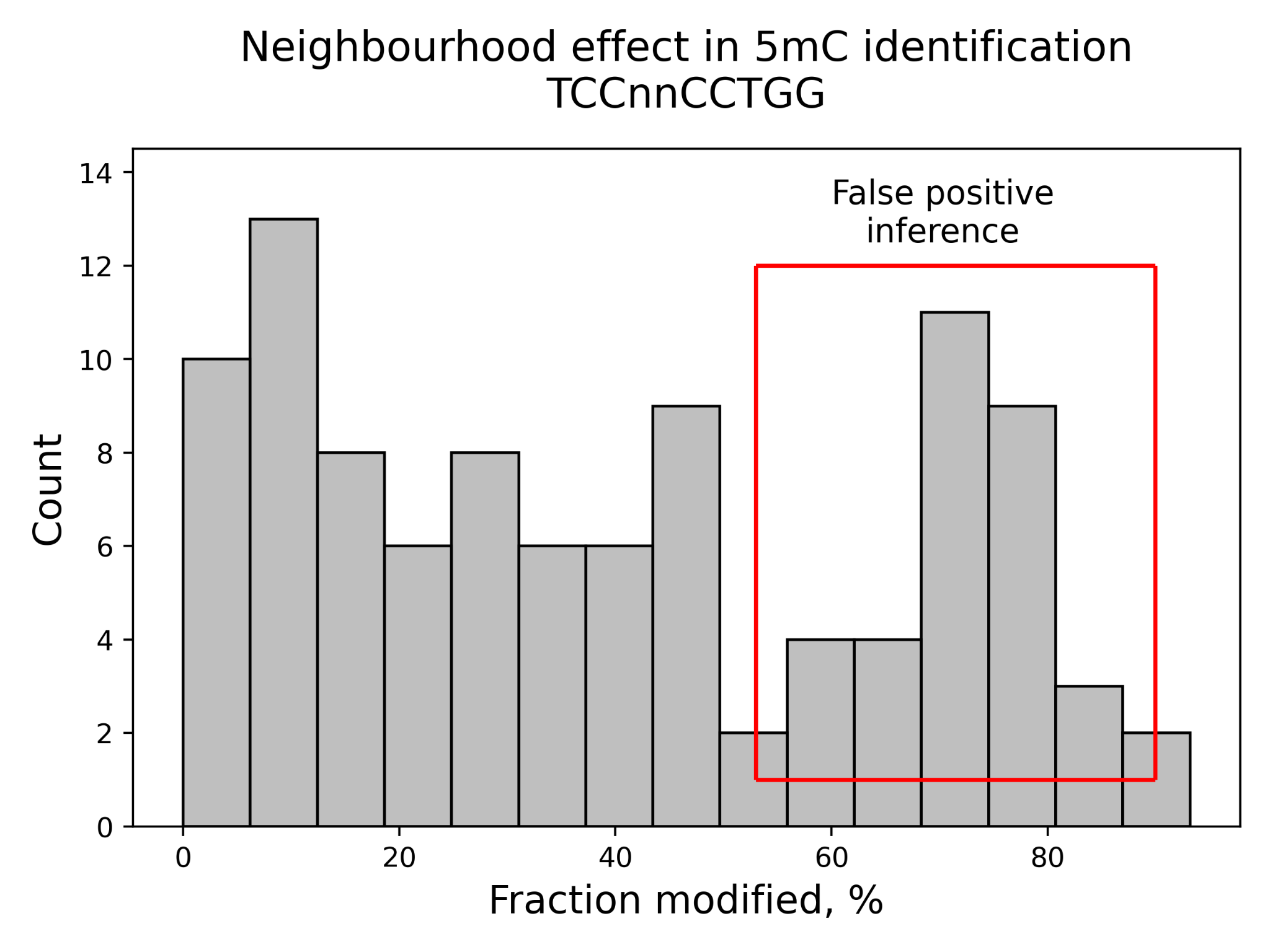

**Supplementary Figure 3.** Demonstration of basecalling neighbourhood effect. This specific histogram represents “fraction modified” values for the site TC**^5m^C**nnCCTGG. This site was not identified by Snappy, but manual observation of the modkit results reveals quite stable methylation of cytosine assigned by the Dorado basecaller in this context. We suppose that high methylation scores in TC**^5m^C**nnCCTGG are most likely an artifact caused by total methylation of a closely located C**^5m^C**TGG motif.

### **Supplementary Data 1. Algorithm Explanation**

#### **Glossary**

***Link* -** the connection of two specific 3-mers in the input sequence. Links represent co-localization of overlapped or closely located 3-mers.

***Length of link*** represents how closely two 3-mers are located in the input sequence. Links with length 1 or 2 represent overlapping 3-mers. links with length 3 represent sequentially located 3-mers, links with length from 4 to 12 represent 3-mers distinguished by 1-9 degenerate positions. Thus, an example of 1-link is a tetramer ACTG (overlapped ACT and CTG 3-mers), an example of 3-link is a hexamer ACTCTG, and an example of 9-link is ACTnnnnnnCTG (the same 3-mers distinguished by any six nucleotides).

***Weight of link*** represents the abundance of this specific link in the input sequence.

***N-position* -** fully degenerate position which could be either A, G, T, or C, and usually is marked as “N” or “n”. In this study, for N-positions we preferably use the literal “.” to simplify writing.

***Modified context (or context)* -** the nucleotide sequence surrounding the modified position. By default it has a length of 31, with 15 bases upstream to the modified base and 15 bases downstream to the modified base.

***Motif variant (or motif)***- the sequence representing potential motif, with fixed length equals the length of *context*. Variants may include degenerate positions, different from N. Typical variants are ………..CATG………., …….CAY……GAT……, ………G.GGHA……, etc.

***Subvariant* -** the sequence and which satisfies a specific *variant,* could have N-positions, but does not have other degenerate positions. For example, ……….GGATC……… is a subvariant for ……….GGHTC……… motif variant but not for ……….GGBTC……… motif variant. Subvariants are used to check correctness and completeness of the motif variant.

***Fraction modified* *value (or frac_mod)* -** for a specific genome position, this value represents percentage (from 0 to 100%) of reads bringing methylation in this position among total occurrences of this position in reads. Similarly, *frac_mod* could be defined for a specific genome site as a percentage of methylated sites among total occurrences of this site in the genome.

***Modality metric*** - the value indicating if a specific motif variant has unimodal *frac_mod* distribution with mode higher than 40%. The modality metric is distributed from 0.0 to 1.0, where values close to 1.0 indicate that this motif variant or subvariant is most likely completely methylated.

#### **Algorithm**

**Required input:** the bed-file resulting from “modkit pileup” command, the reference genome

1) First, the algorithm constructs a graph structure that represents the frequency (weights) of *links* between all canonical 3-mers presented in the input sequence data. In the graph structure, each node represents 3-mer, and each node pair is connected by 12 weighted and labeled edges, where labels represent different *link lengths*, and edge *weights* represent the frequency of specific 3-mer links in the input data. Therefore, the total number of nodes in the graph equals 4^3^, and the total number of of edges equals 4^3^ * 4^3^ (all possible 3-mer pairs) * 12 (all possible link lengths) = 49152, so in computational terms such a structure is quite compact.	This graph structure is built for the target sequences (i.e. modified contexts) and for the control sequences (usually the whole genome) independently.
2) Next, using Chi-square test, the algorithm searches for the edge in the graph, which is most significantly over-weighted in the target graph compared to the control. This edge represents a short nucleotide motif, which next will be used as an anchor motif. Since 1- and 2-links represent overlapped 3-mers, there could be from 4 to 6 canonical bases in the anchor motif. Examples of anchor motifs are GATC, AGTACT, and ATG…….CGT. In the end of this step, the algorithm identifies the preferred position of the chosen anchor in the modified contexts with length of 31 and converts it to corresponding 31-mer (for example n_13_GATCn_14_, n_14_AGTACTn_11_, or n_14_ATG…..CGTn_6_), constructing an initial motif variant.
3) Next, using ModKit results, the algorithm collects *frac_mod* values for all the modified contexts satisfying the initial motif variant and checks if they provide unimodal distribution with a modality metric greater than 0.9. If they are, the algorithm goes to the next step. Otherwise, it tries to extend the motif, iteratively changing N-positions adjacent to canonical bases to A, G, T, or C. Based on the modality metric, the algorithm chooses the best extension variant, modifies the motif, and repeats the procedure if necessary. When an extended motif provides the required modality metric value, the extending process stops, and the algorithm goes to the next step. If the algorithm cannot find an optimal anchor extension, it terminates and returns the result.
4) In this step, the inferred motif is supposed to be fully modified but incomplete, i.e. it could be just a subvariant of the actual methylation motif (for example, the initial motif is GAGTC while the actual motif is GANTC). To fix that, the algorithm iteratively changes each non-N-position of the motif to a more degenerate base (S, Y, B, H, N, etc). If after the change modality metric remains to be higher than 0.9, and the metric of all subvariants of the new variant is higher than 0.9, the algorithm applies this change to the motif and repeats the procedure. This procedure stops, when the algorithm could not find a satisfactory change.
5) The resulting sequence is considered as a fully methylated and complete motif. The Chi-square statistics value is computed for the inferred motif. If the confidence metric for the current motif is greater than 100, the algorithm adds the motif to the final results. Regardless of the confidence value, the modified contexts satisfying the motif are filtered from the input target sequences. If the total number of modified contexts after filtering is less than 20, the algorithm terminates its work and returns the result. Otherwise, it uses the filtered modified contexts to construct a new target graph of 3-mer links and goes to step 2 trying to find another motif.

**Output:** the list of identified methylation sites sorted by confidence values.

### **Supplementary Data 2. Snappy’s rules**

1) Snappy does not use any heuristics. Snappy does not know what methylation motifs are, and how they should look like.
2) Snappy deliberately does not make assumptions if a context is modified or not when there are no such contexts in the input genome. It is why Snappy could return non-complete motifs in some cases.
3) Snappy deliberately does not make assumptions if there could be neighbourhood effects. (Snappy just doesn’t know what the neighbouring effect is.) It returns all motifs, which satisfy its logic.
4) Similarly, Snappy does not know about the effects caused by homopolymeric regions. It is a problem of a basecaller, not Snappy.
5) Snappy deliberately does not try to filter motifs with bimodal distribution of frac_mod, if there are no modes close to zero in the distribution. Snappy is simple, and thinks that non-modified bases should have values of frac_mod close to zero. Users can filter such cases by themselves, using visualization provided by Snappy.
6) In general, Snappy does exactly what it is asked to do. It searches for motifs that are over-represented in the modified contexts, and that provide frac_mod distributions without zero-modes. Any interpretation of the results is the responsibility of the user.
